# EBV INFECTION OUTCOMES DETERMINED BY MONOCYTE AND T_REG_-DRIVEN IMMUNE DYNAMICS IN AN EX VIVO PBMC MODEL

**DOI:** 10.1101/2025.11.20.689441

**Authors:** Leena Yoon, Lauren N. MacMullen, Leonardo Josué Castro Muñoz, Alina Gu, Jamie Bregman, Mary Campion, Avi Srivastava, Rena R. Xian, Richard F. Ambinder, Andrew Kossenkov, Samantha S. Soldan, Paul M. Lieberman

## Abstract

Epstein-Barr virus (EBV) infects >95% of the adult population with diverse outcomes ranging from benign latency to cancers and autoimmune diseases. Immunological control of EBV infection is known to be an important determinant of EBV infection outcomes. However, species-specific viral tropism and limited infection models have impeded mechanistic insights into early host–immune control of EBV infection. Here, we use *ex vivo* infection of peripheral blood mononuclear cells (PBMCs), rather than routinely used B cell enriched culture systems, to study immune and viral dynamics during primary EBV infection. We combined bulk RNA sequencing, EBV transcript enrichment, and flow cytometry to characterize cellular responses across Days 1, 7–8, and 14 post-infection. Early infection triggered a monocyte-specific antiviral response marked by changes in the expression of genes associated with lipid metabolism (LIPA, lysosomal acid lipase) and chemotaxis (CCR1 and CCR2). Inhibitors of LIPA increased EBV titers during primary infection, indicating that LIPA is part of an early monocyte-driven antiviral response. At later timepoints post-infection, donor-dependent variability in lymphoblastoid cell line (LCL) outgrowth was associated with divergent immune states. Donors that failed to generate LCLs demonstrated increased frequencies of CD8^+^ T cells and reduced numbers of regulatory T cells (CD4⁺CD25⁺FOXP3⁺). EBV transcriptomics revealed that LCL-failed donors exhibited elevated early lytic gene expression but did not establish a type III latency program. Our findings suggest that individual variations in immune cell composition and gene expression may account for differences in the immune response to EBV. These findings define temporal immune and viral signatures that predict transformation outcome and highlight intact PBMCs as a tractable model to study EBV pathogenesis in a genetically diverse, human-specific context.

**AUTHOR SUMMARY:** Individual variation in response to Epstein-Barr virus (EBV) infection can lead to diverse pathogenic outcomes, ranging from cancers to autoimmune disease. To study this variation, we analyzed immune cell response and viral dynamics during the ex vivo primary EBV infection of peripheral blood mononuclear cells (PBMCs) from donors that either fail or succeed to generate lymphoblastoid cell lines (LCLs). Flow cytometry and RNA-seq revealed a rapid monocyte-specific antiviral response among all donors marked by genes associated with lipid metabolism (LIPA) and chemotaxis (CCR1 and CCR2). LIPA inhibition increased EBV titers during primary infection, demonstrating a functional antiviral role. At later timepoints, donor-specific differences in CD8^+^ T cells and Treg subsets, along with EBV gene expression, were correlated with successful LCL outgrowth. Treatment with the Treg-depleting antibody RG6292 suppressed viral transformation in donors that otherwise supported LCL outgrowth, confirming a functional role for Tregs in shaping early EBV infection outcomes. Viral transcript enrichment-seq revealed an upregulation of early lytic and failure to sustain latent gene expression correlating with failure to generate LCL. These findings highlight intact PBMCs as a tractable model to study EBV viral-host interaction in a genetically diverse, human-specific context, and that Tregs play a key determining role in viral transformation.

## INTRODUCTION

Epstein-Barr virus (EBV), also known as human herpesvirus 4 (HHV4), was identified as the first human tumor virus, but is now known to infect over 95% of the global population [1–4]. While most EBV infections are subclinical, EBV is also known to be a causal agent in various lymphoid and epithelial cell malignancies, and more recent studies implicate EBV as a causal factor in autoimmune disease, especially multiple sclerosis [1, 5, 6]. In most healthy adults, EBV establishes a long-term latent infection in memory B-cells with periodic reactivation and chronic shedding in the oropharynx. In some individuals, EBV primary infection can cause a hyper immune response resulting in infectious mononucleosis (IM), characterized by an expansion of infected B-cells and reactive T-cells[7, 8]. Severe IM is a major risk factor for developing multiple sclerosis (MS) [9]. On the other hand, immunosuppression and inherited immunodeficiencies are major risk factors for EBV-associated malignancy [10]. The factors that determine benign verse pathogenic EBV infection outcomes are not completely known, but the balance of infection efficiency and immune control are thought to be major determinants.

EBV is well known for its ability to efficiently transform resting B-cells into continuously proliferating lymphoblastoid cell lines (LCLs) [11, 12]. LCL formation requires both the transformation of B-cells as well as the elimination of T-cells that would otherwise control EBV infected cell proliferation. Most experimental models of EBV immortalization have utilized *ex vivo* infection of purified B cells [13–18]. The studies have provided detailed understanding of the molecular process through which EBV can efficiently immortalize purified human B cells *in vitro*, including mechanisms of viral entry, gene regulation, cell proliferation, anti-apoptosis, and cell-cell signaling [19–23]. EBV infection of purified B-cells typically results in efficient transformation of naïve B-cells into stable lymphoblastoid cell lines (LCLs) within the first 4 weeks under optimal conditions [13]. While there is important heterogeneity in B-cell response to EBV infection [24], most human donors successfully generate LCLs from purified B-cells. However, these purified B-cell models fail to capture immune cell interactions which can also influence B-cell immortalization and immune responses to EBV primary infection, highlighting the need for broader cellular models to better elucidate EBV-host interactions [25–27].

Several experimental models capture some of the complexity of immune interactions that determine EBV infection outcomes [28–30]. Because EBV is highly human species specific, only a few animal models have been tractable for study of human disease. The humanized mouse models provide valuable insights into the immune control of lymphoproliferative disease and autoimmune outcomes, but this model is limited by technical and resource challenges [30, 31]. EBV *ex vivo* infection of purified primary B-cells has been studied extensively, including at the single cell RNA-seq level to reveal mechanisms of viral induced B-cell immortalization and diversity of B-cell responses to EBV infection [17, 32–34]. While these studies offer valuable insights into B-cell heterogeneity and latent transformation, they have not resolved early viral transcription due to low infection rates, limited capture efficiency, and restricted sampling of productively infected cells. In addition, purified B cell models lack immune cell diversity, which precludes assessment of non–B cell responses and intercellular signaling. These limitations underscore the need for experimental systems that preserve a more complete immune repertoire and allow temporal resolution of both host and viral dynamics during primary EBV infection. Human peripheral blood mononuclear cells (PBMCs) provide a physiologically relevant *ex vivo* model that preserves both immune cell and genetic diversity, and an opportunity to investigate molecular details of primary infection in the context of a complex immune system. This system enables the simultaneous study of direct viral infection and immune-mediated responses within a native cellular context.

In this study, we used human PBMCs to characterize the cellular immune response and transcriptional dynamics of early EBV infection at Day 1, 5, 7–8, and 14 post-infection. We combined immunophenotyping and bulk RNA-seq to characterize the sequential immune response to EBV infection. We identified an early response (day1) dominated by monocytes, where a marked upregulation of inflammatory and lipid metabolism genes. At later time points (days 5-14), responses varied depending on donor-specific infection outcomes. Donors that failed to generate LCL showed reduced CD4⁺CD25⁺FOXP3⁺ Tregs and different cytotoxic responses, suggesting that lack of T reg suppression restricts transformation and latency establishment. By enriching for EBV transcripts, we mapped viral gene expression dynamics and linked lytic-to-latent transition patterns with host immune control. These findings establish PBMCs as a robust platform to study host–virus interactions and provide mechanistic insight into how individual variation in immune response shapes EBV infection outcomes relevant to persistence, transformation and autoimmunity.

## RESULTS

### *Ex Vivo* EBV Infection of PBMCs Reveals Time-Dependent Immune Remodeling

To model early immune dynamics following EBV exposure, we infected human PBMCs *ex vivo* with MUTU1 strain of EBV and monitored cellular responses at different time-points (**Figure 1A**). All donors used in this study were seropositive for EBV, ensuring prior immune priming, a critical factor for interpreting transformation outcomes (**Supplementary Table S1**). By later stages, we observed striking inter-donor variability in the ability to generate lymphoblastoid cell lines (LCLs): while some donors reproducibly failed to establish LCLs, others made LCLs with success rates exceeding 90% (**Figure 1B, Supplementary Figure S1**). All donors and infections could generate LCLs if T-cell immunosuppressing drug cyclosporin was added during the infection (**Supplementary Table S2**). To assess early infection, we measured the EBV surface protein gp350 and cell activation marker CD23 by flow cytometry (**Fig. 1C**). These markers revealed distinct infection trajectories: LCL-made donors exhibited a sustained increase in gp350⁺CD23⁺ cells over time, while LCL-failed donors showed an initial rise by Day 7 followed by a sharp decline. To determine how these different outcomes were associated with early immune compositional shifts, we performed flow cytometric immunophenotyping coupled with t-distributed stochastic neighbor embedding (t-SNE) visualization. Gating strategies for immune cell phenotyping are shown in **Supplemental Fig. S2, S3**. This analysis revealed pronounced shifts in immune cell populations, including a decline in B cells and expansion of T-cells across all donors by Day 7. However, in LCL-forming cultures, B cell frequencies expanded substantially by Day 21, reaching approximately 40% of the live cell population, and continued to increase by Day 28 with B cells representing most viable cells. In contrast, LCL-failed donors exhibited continuous B cell depletion, accompanied by changes in T-cell subsets with progressive loss of total cell population viability, dropping down to 10–15% by Day 21, with minimal recovery thereafter (**Figure 1C**). These findings demonstrate that EBV infection of PBMC *ex vivo* captures human donor-specific immunological diversity and plasticity.

**Figure 1.**
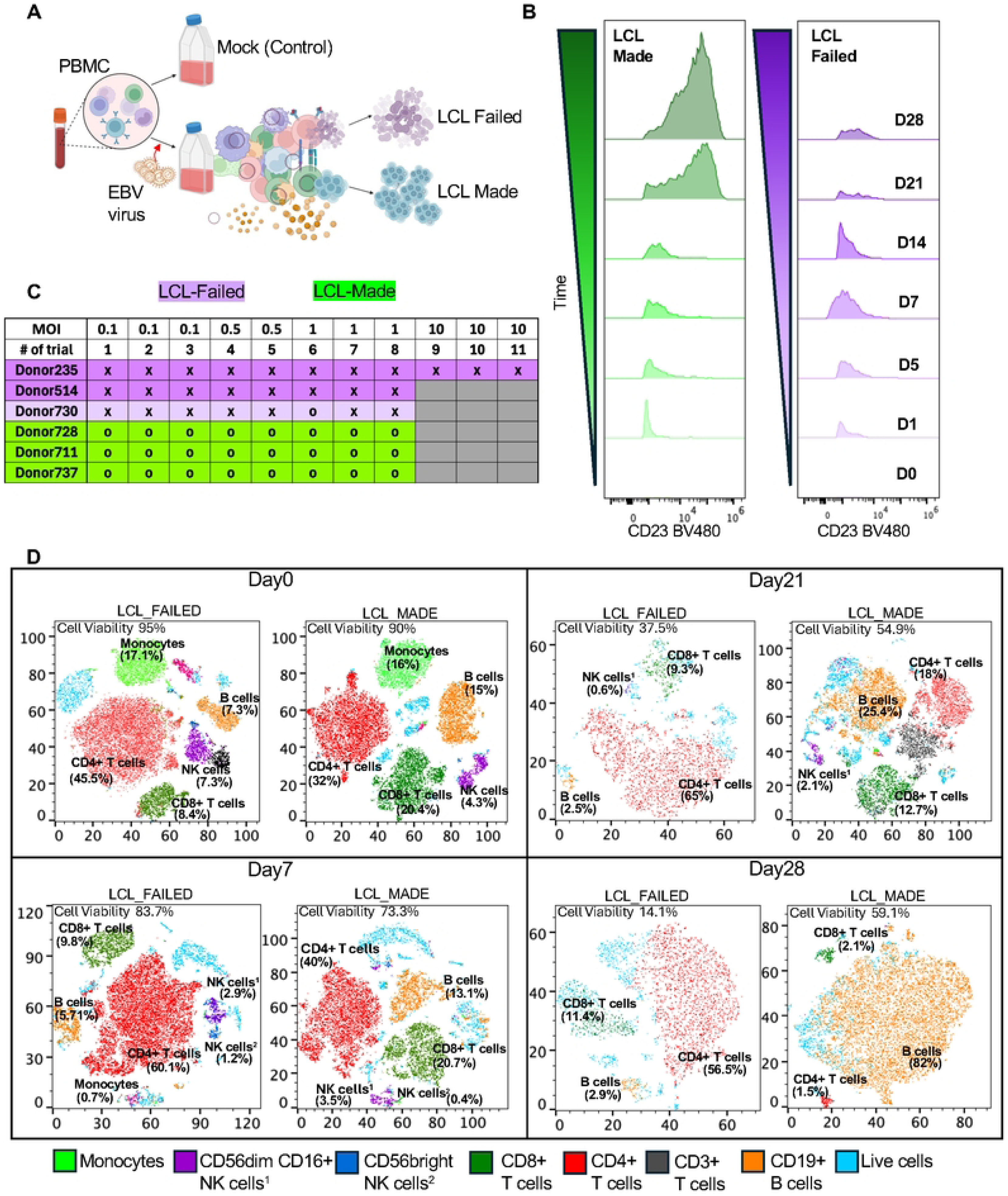
*Ex Vivo* EBV Infection of PBMCs Reveals Donor-Dependent Differences in B Cell Transformation Potential. (A) Schematic of the *ex vivo* infection model. Human PBMCs were infected with the Mutu I strain of EBV at MOI = 1 and harvested at indicated time points. Donors were later classified as LCL-made or LCL-failed based on transformation outcome. (B) Representative flow cytometry plots showing gp350⁺CD23⁺ cells across time points in LCL-made and LCL-failed donors. LCL-made donors exhibited a sustained increase in infected B cells, while LCL-failed donors showed minimal or transient induction. (C) LCL outcome classification was based on repeated infection assays (>3 per donor and MOI), with LCL-made (o) or LCL-failed (x) groups defined by >90% success or failure rates, respectively. (D) t-SNE plots showing longitudinal shifts in immune cell composition and viability in LCL-made and LCL-failed donors at Day 0, 7, 21, and 28. Flow cytometry determined percentages of cell type are shown for Monocytes (green), CD56^dim CD16^+^ NK cells (purple), CD56^bright NK cells (blue), CD8^+^ T cells (green), CD4^+^ T cells (red), CD3^+^ T cells (black), CD19^+^ B cells (orange), live cells (light blue).

### EBV Infection Induces Time-Resolved Transcriptomic Changes in Human PBMCs

To further characterize the immune dynamics following EBV infection, we performed bulk mRNA sequencing on the same samples. Principal component analysis (PCA) revealed a clear temporal separation of samples among Day 1, Day 5–8, and Day 14, indicating a progressive and coordinated shift in the global immune transcriptome (**Figure 2A**). To capture broad transcriptional effects of EBV infection, we performed Gene Ontology (GO) enrichment analysis across all donors and time points. Enriched pathways included viral response programs such as regulation of viral entry into host cell and regulation of viral genome replication, consistent with active infection. Immune regulatory terms such as positive regulation of T cell cytokine production, positive regulation of defense response to virus, and cytokine-mediated signaling were also enriched. Pathways related to cell proliferation, including stem cell proliferation and T cell activation, were among the top-ranked categories. In parallel, we observed enrichment of stress response pathways, including MAPK cascade and neuroinflammatory regulation, suggesting broader cellular remodeling. These results indicate that EBV infection induces coordinated immune, stress, and viral programs across PBMCs (**Figure 2B**). A heatmap of differentially expressed genes revealed a uniform response at Day 1, but then a striking divergence among individual donor profiles at later time points (**Figure 2C**).

**Figure 2.**
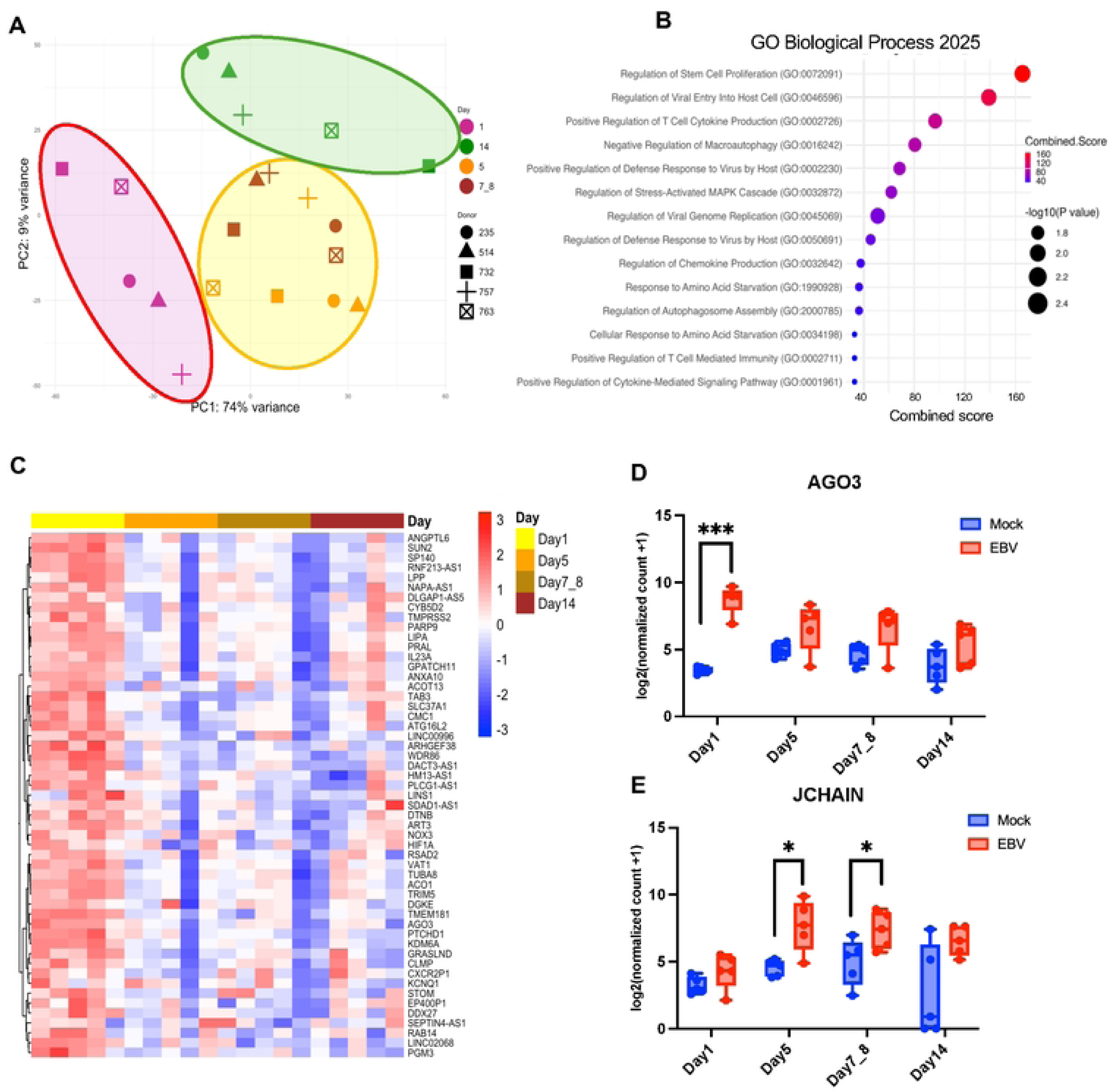
Bulk Transcriptomic Profiling Reveals Time-Dependent Gene Expression Changes Following EBV Infection. (A) Principal component analysis (PCA) of bulk RNA-seq data for samples across Days 1, 5, 7–8, and 14 post-infection (color icons), and donors (shaped icons). Group circle indicates PCA separation along time points 1 (purple), 5-8 (yellow), and 14 (green) days after infection. (B) Gene ontology (GO) enrichment analysis of differentially expressed genes (FDR < 0.05) across all timepoints highlights key biological processes modulated by EBV infection. (C) Heatmap of top differentially expressed genes clustered by days post-infection. List of top 50 most significantly regulated genes indicated to the right. (D, E) Representative genes illustrating distinct temporal expression patterns following EBV infection for AGO3 (panel D) and JCHAIN (panel E). Data are presented as mean ± s.d. from n=5 donors. Statistical significance was determined using a paired two-tailed Student’s t-test.

Transcriptomic analysis revealed distinct temporal patterns of host gene regulation following EBV infection. Some genes, such as AGO3, showed transient induction restricted to Day 1, reflecting an acute antiviral or stress-associated response (**Figure 2D**). Others, including JCHAIN, a polypeptide that regulates the formation of multimeric antibodies, were selectively modulated at later timepoints, indicating delayed or secondary immune remodeling (**Figure 2E**). In contrast, genes like HIF1A and KLRF1 were consistently upregulated across all timepoints in infected samples (**Supplemental Figure S4**), suggesting sustained involvement in metabolic adaptation and innate cytotoxic responses. These data indicate that an early uniform response to EBV occurs across all donors, while at later timepoints after Day 1 transcriptional response kinetics vary dramatically among donors, reflect a mult-phase immune-metabolic reprogramming of immune cells that appear to segregate with infection outcomes.

### Early EBV Infection Triggers Immune Activation

Analysis of RNA-seq of PBMCs infected ex vivo with the EBV revealed an early host response by Day 1 post-infection, with over 800 differentially expressed genes (false discovery rate [FDR] < 0.05). A volcano plot (**Figure 3A**) illustrated the global transcriptional shift, while a curated heatmap of top-ranked genes (**Figure 3B**) revealed consistent induction patterns across biological replicates. Among the most strongly upregulated genes were LIPA (lysosomal acid lipase), KDM6A (a chromatin-modifying enzyme linked to immune regulation)[35], and RSAD2 (also known as Viperin), an interferon-stimulated gene (ISG) with known antiviral activities[36]. These findings suggest that early EBV exposure elicits a coordinated antiviral, metabolic, and epigenetic transcriptional program in peripheral immune cells.

**Figure 3.**
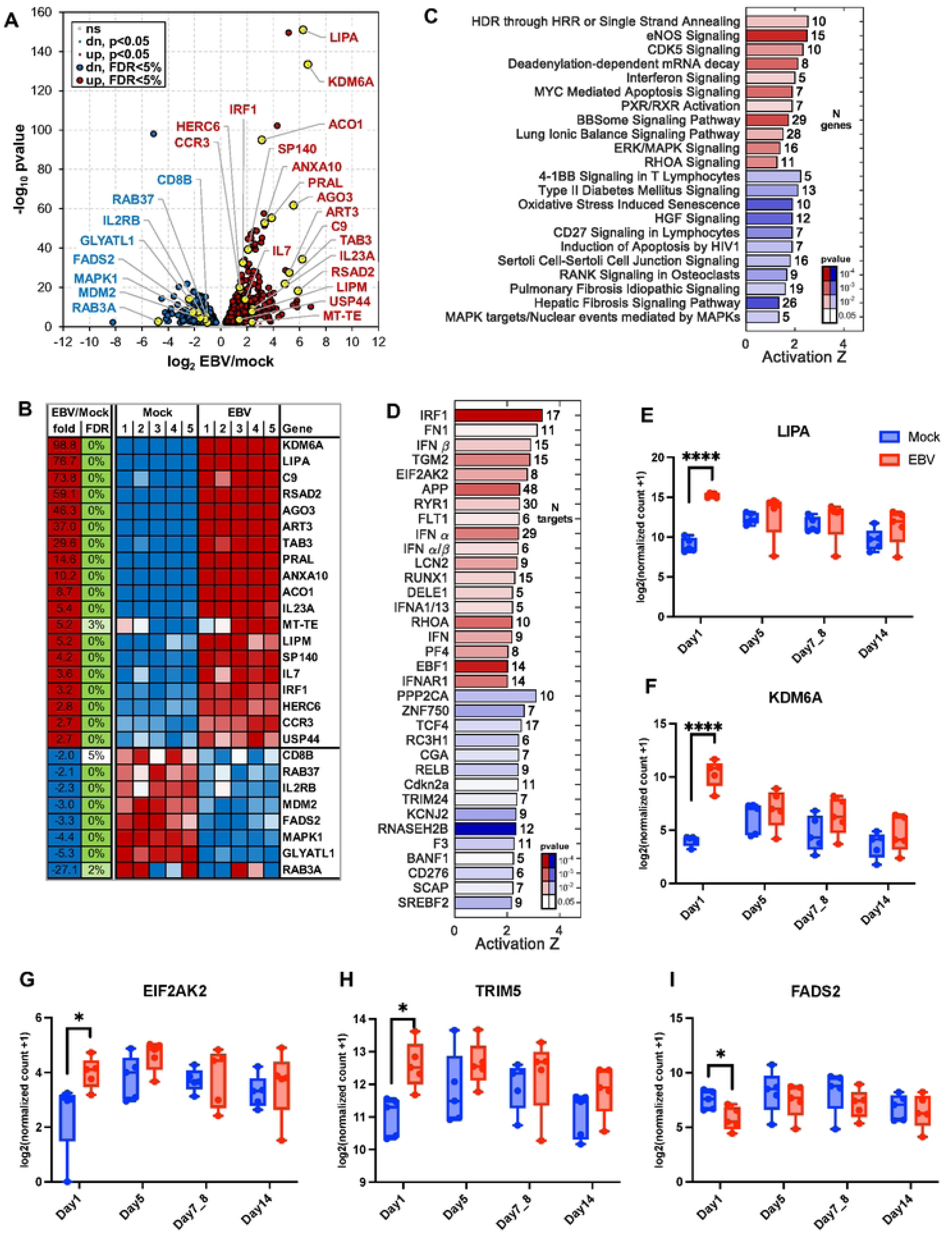
Early Transcriptional Responses to EBV Infection at Day 1 Post-Infection. (A) Volcano plot showing differentially expressed genes at Day 1 post-infection compared to mock. (B) Heatmap of selected differentially expressed genes with FDR < 0.05. (C) Gene set enrichment analysis highlights the top 22 pathways significantly affected at Day 1 (activation Z-score > 1, p < 0.005). Red indicates activated pathways; blue indicates inhibited pathways. (D) Top 34 predicted upstream regulators with significant activation or inhibition (Z > 2, p < 0.05). MicroRNAs and chemical/drug regulators were excluded. (E–I) Representative genes showing differential expression at Day 1: (E) LIPA, (F) KDM6A, (G) EIF2AK2, (H) TRIM5, (I) FADS2. Data are mean ± s.d. from n = 5 donors. Statistical significance was assessed using a paired two-tailed Student’s t-test.

Pathway enrichment analysis identified early engagement of immune, stress, and metabolic pathways, including type I interferon signaling, oxidative stress-induced senescence, MYC-mediated apoptosis, and ERK/MAPK pathways (**Figure 3C**). Enrichment of CD27 and 4-1BB pathways in T cells further supports early activation of co-stimulatory immune signals. To identify upstream regulators, we used Ingenuity Pathway Analysis (IPA), which identified IRF1, IFN-β, and EIF2AK2 (PKR) as top predicted activators (**Figure 3D**), consistent with an environment primed for antiviral defense and oxidative stress responses. To delineate early specific responses, we identified genes with altered gene expression only at Day 1. Genes such as LIPA, KDM6A, EIF2AK2, and TRIM5 were upregulated in EBV-infected PBMCs, while FADS2, a fatty acid desaturase, was downregulated (**Figures 3E–I**). These findings show that EBV exposure initiates a multilayered transcriptional response that involves innate immune activation, interferon signaling, oxidative stress, and lipid metabolic remodeling, and point to downstream immune modulation and cellular transformation.

### EBV Induces Monocyte-Specific LIPA Expression and Alters Chemokine Receptor Profiles

To better understand the upregulation of LIPA, we performed flow cytometry to identify the cell type where this gene and protein were most upregulated during EBV infection of PBMC. Flow cytometry revealed that LIPA expression is highest in monocytes compared to B and T cells and increases specifically in monocytes after EBV exposure (**Fig. 4A, Supplementary figure S5A**). These findings were further validated by qPCR analysis, which confirmed transcriptional upregulation of LIPA in EBV-infected PBMCs (**Supplementary Figure S5B**). LIPA encodes lysosomal acid lipase (LAL), which hydrolyzes cholesterol esters and triglycerides into free fatty acids and cholesterol. To assess LIPA function, we inhibited LAL using Lalistat2 and observed a significant increase in EBV viral load compared to untreated controls (**Fig. 4B**), which supports a role for LIPA-driven lipid metabolism in early antiviral defense. In addition, EBV infection triggered a monocyte-specific immune response detectable at Day 1, based on expression profiles in CD14⁺CD64⁺ monocytes. Expression of the monocyte-associated chemokine receptors CCR1 and CCR2 was significantly reduced in infected samples to mock controls (**Fig. 4C–E**), suggesting that EBV exposure reshapes monocyte migration potential and inflammatory phenotype.

**Figure 4.**
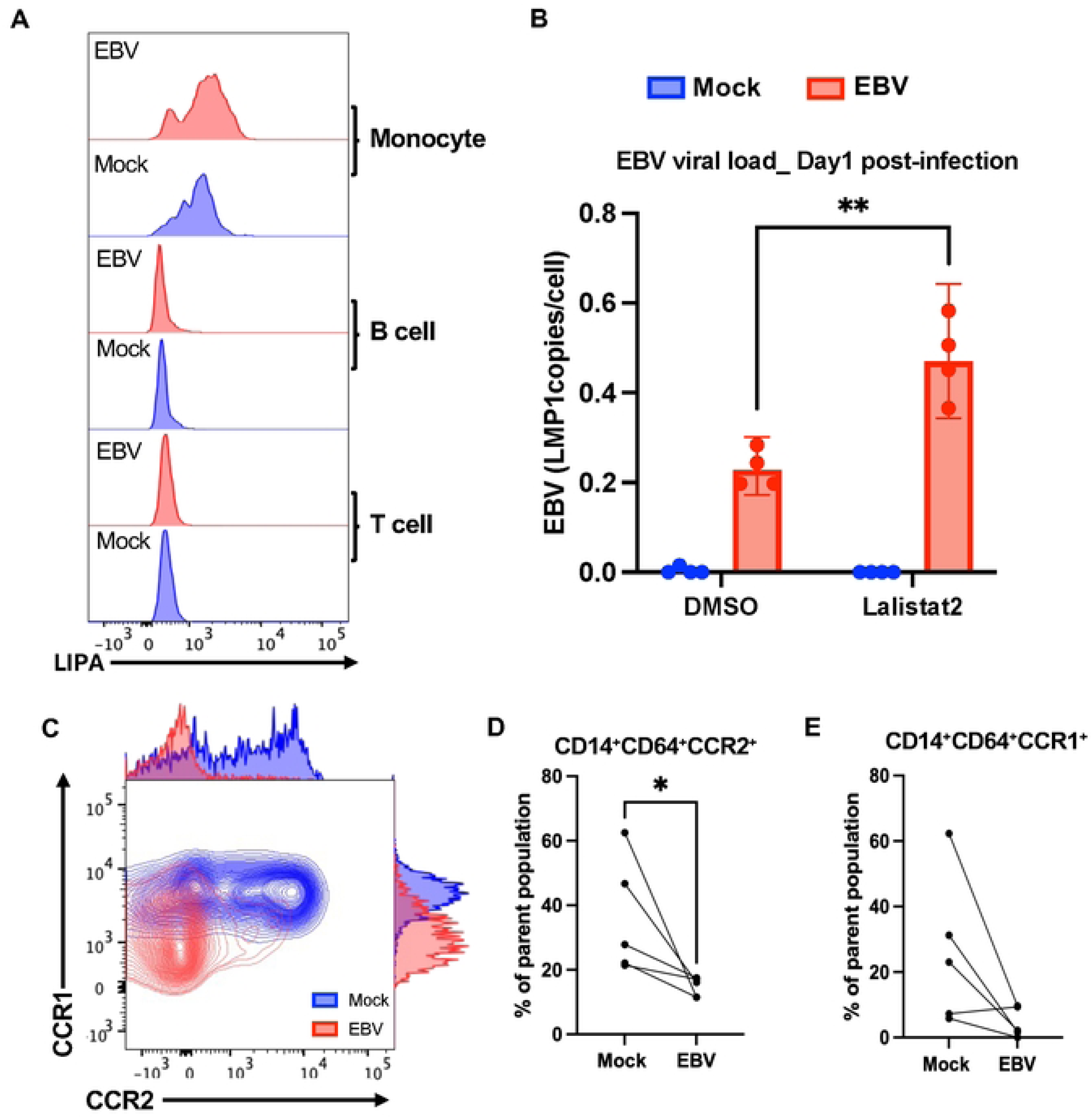
LIPA and Chemokine Receptor Remodeling Mark Monocyte Activation in Early EBV Infection. (A) Flow cytometry analysis of LIPA expression across immune subsets at Day 1 post-infection. LIPA levels were elevated in EBV-infected (red) versus mock-treated (blue) samples, with induction restricted to CD14⁺ monocytes. No significant changes were observed in B or T cells. (B) EBV DNA copy number was measured by ddPCR after pharmacological inhibition of lysosomal acid lipase using Lalistat2. Data are mean ± s.d. from n = 4 donors. Statistical significance was assessed using a paired two-tailed Student’s t-test. (C) Representative flow cytometry plots show reduced expression of chemokine receptors CCR1 and CCR2 in monocytes following EBV infection. (D–E) Quantification of CCR1 and CCR2 expression in CD14⁺CD64⁺ monocytes show significant downregulation upon EBV exposure. Data are mean ± s.d. from n = 5 donors. Statistical significance was assessed using a paired two-tailed Student’s t-test.

### Reduced Treg and Absence of T Cell Exhaustion Characterize Donors That Fail to Support LCL Outgrowth

From Day 7–8 post-infection, distinct donor-dependent variability in B cell immortalization emerged. Some individuals consistently supported EBV-driven lymphoblastoid cell line (LCL) outgrowth, while others failed to generate LCLs despite repeated attempts (**Figure 1C, Supplementary Table S2**). Each donor was tested in more than 10 independent infection assays under standardized conditions using the same EBV strain and culture setting. Despite variation in multiplicity of infection (MOI) and repeated trials, transformation outcomes remained consistent. Donors with >85% success or failure rates were classified as LCL-made or LCL-failed, respectively, and those with intermediate outcomes were excluded from downstream analyses (**Supplementary Table S2**).

Transcriptomic profiling revealed marked differences between the two groups (**Fig. 5A, Supplemental Figure S6**), which suggest the presence of an intrinsic immune response signature associated with LCL outcome. While both Day 7 and Day 14 samples showed separation between groups, we focused on Day 14 for downstream analyses due to stronger transcriptional divergence and clearer within-group separation, particularly in LCL-made donors across mock and EBV conditions. Volcano plot and hierarchical clustering identified the top differentially expressed genes distinguishing between LCL-made (green) and LCL-failed (purple) donors (**Figure 5B, C, and Supplemental Figure S7A**). Genes such as CCL13, STAT5B and CCR3 were consistently downregulated in LCL-failed donors. In contrast, genes including CD44, NFRKB, ROCK2, and ACAT1 were upregulated in the LCL-failed group (**Supplementary Figure S7**). These differentially expressed genes reflected multiple immune modules: immune signaling and cytokine regulation (STAT5B), chemotaxis (CCL13, CCR3), and metabolic adaptation (MDH1, METTL9). Pathway analysis highlighted activation of IL-6–STAT5B signaling, leptin-triggered cytokine production, ROS-linked neutrophil cytotoxicity, and chemokine receptor regulation (**Figure 5D**).

**Figure 5.**
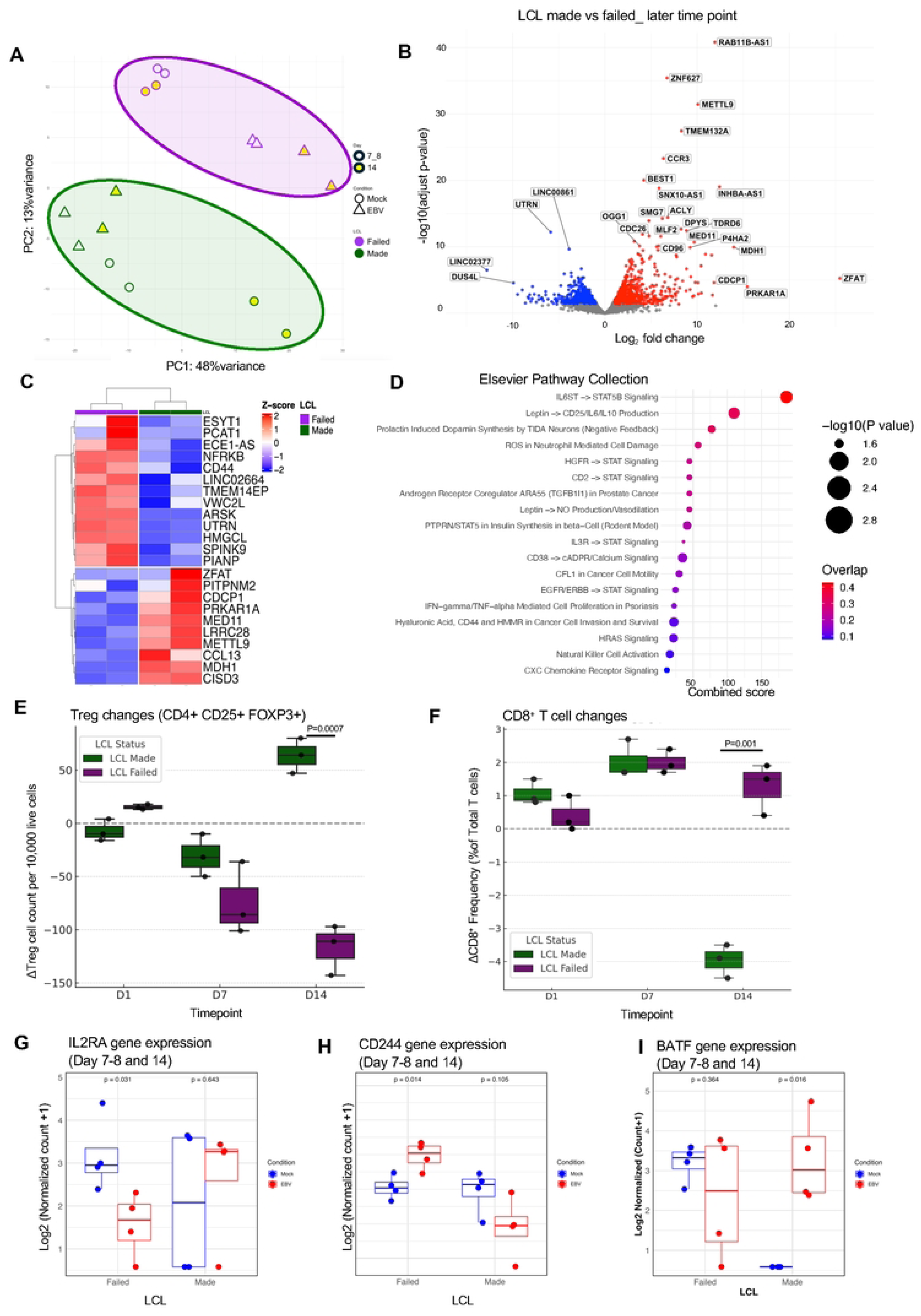
Distinct Transcriptional Signatures at Day 7–8 Define LCL Transformation Outcome. (A) Principal component analysis (PCA) of global transcriptomes from EBV-infected PBMCs at Day 7–8 and Day14 (yellow). Samples stratify according to LCL outcome, separating LCL-made (green) from LCL-failed (purple) donors. (B) Volcano plot showing differentially expressed genes between LCL-made and LCL-failed donors (FDR < 0.05). Red and blue denote significantly up-and downregulated genes in LCL-failed donors, respectively. (C) Heatmap of the top differentially expressed genes showing unsupervised hierarchical clustering at Day 14. (D) Pathway enrichment analysis of differentially expressed genes using the Elsevier Pathway Collection. The x-axis shows the Combined Score, calculated as the log-transformed p-value multiplied by the z-score of the deviation from expected rank. (E) Frequencies of regulatory T cells (CD4⁺CD25⁺FOXP3⁺) measured by flow cytometry (mean ± SEM; p < 0.05, ns = not significant; unpaired two-sided Student’s t-test (n=3)) (F) Frequencies of CD8⁺ T cells measured by flow cytometry (mean ± SEM; p < 0.05, ns = not significant; unpaired two-sided Student’s t-test (n=3)) (G) RNA-seq expression levels of IL2RA at Day 7-8 and 14 (H) RNA-seq expression levels of CD244 in PBMCs at Day 7-8 and 14, stratified by LCL outcome. Statistical significance was assessed using a paired two-sided Student’s t-test. (I) RNA-seq expression levels of BATF in PBMCs at Day 7-8 and 14, stratified by LCL outcome. Statistical significance was assessed using a paired two-sided Student’s t-test.

These transcriptional shifts also suggest changes in immune cell composition. From Day 7, flow cytometry showed that regulatory T cell (Treg) frequencies dropped in LCL-failed donors but remained stable in LCL-made donors (**Figure 5E**). Tregs were defined as CD3⁺CD4⁺CD25⁺FOXP3⁺, and frequency changes were calculated as change in Treg cell number in EBV infected compared to mock. In contrast to Treg contraction, we observed a significant reduction of total CD8⁺ T cells in LCL-made donor PBMCs, while CD8⁺ T cells showed an increase in the LCL-failed group (**Figure 5F**). At the transcriptomic level, Treg changes aligned with decreased IL2RA expression in LCL-failed donors (**Figure 5G**), a gene essential for Treg survival and homeostasis [37]. Similarly, CD8⁺ T cell changes corresponded with differential expression of CD244, a surface receptor associated with CD8⁺ T cell activation and early exhaustion [38, 39]. CD244 expression was significantly elevated in the LCL-failed group (p = 0.014), whereas it was reduced or unchanged in LCL-made donors (p = 0.105) (**Figure 5H**). Markers of T cell exhaustion exhibited distinct expression patterns across the groups (**Figure 5I; Supplementary Figure S8**). LCL-made group showed increased expression of exhaustion-related genes, including BATF [40], HAVCR2 (TIM-3) [41], and ENTPD1 (CD39) [42], while these markers were either downregulated or unchanged in LCL-failed donors. Notably, TOX [43] and SLAMF6 [44], key markers of terminal and progenitor exhaustion, respectively, were decreased in LCL-failed donors, suggesting the absence of both terminally exhausted and stem-like CD8⁺ T cell subsets in this group.

Together, these data suggest that Tregs may support EBV persistence in LCL-made donors by maintaining an exhausted T cell environment, while the expansion of activated, non-exhausted CD8⁺ T cells in LCL-failed donors may reflect a more robust antiviral response that ultimately interferes with B cell transformation.

### Donor-specific Treg dynamics control early EBV infection and LCL outcome

To validate immune subset shifts observed in bulk RNA-seq, we performed single-cell RNA-seq on PBMCs at day 7 post-infection. UMAP plots revealed clear separation of major immune lineages, including Tregs, B cells, CD4⁺ and CD8⁺ T cells, NK cells, and myeloid cells (**Figure 6A and B**). To quantify infection-induced changes, we calculated the difference in cell type proportions between EBV-infected and mock samples (Δ = EBV – Mock). Stratification by LCL outcome revealed distinct trends: LCL-failed donors showed reduced B cells, myeloid cells, and NK cells, and increased CD8⁺ T cells, while LCL-made donors exhibited increased B cells and reduced CD8⁺ and CD4+ T cells (**Figure 6C**). These patterns were not observed when grouping by infection status alone (**Figure 6B**), highlighting donor outcome as a key variable.

**Figure 6.**
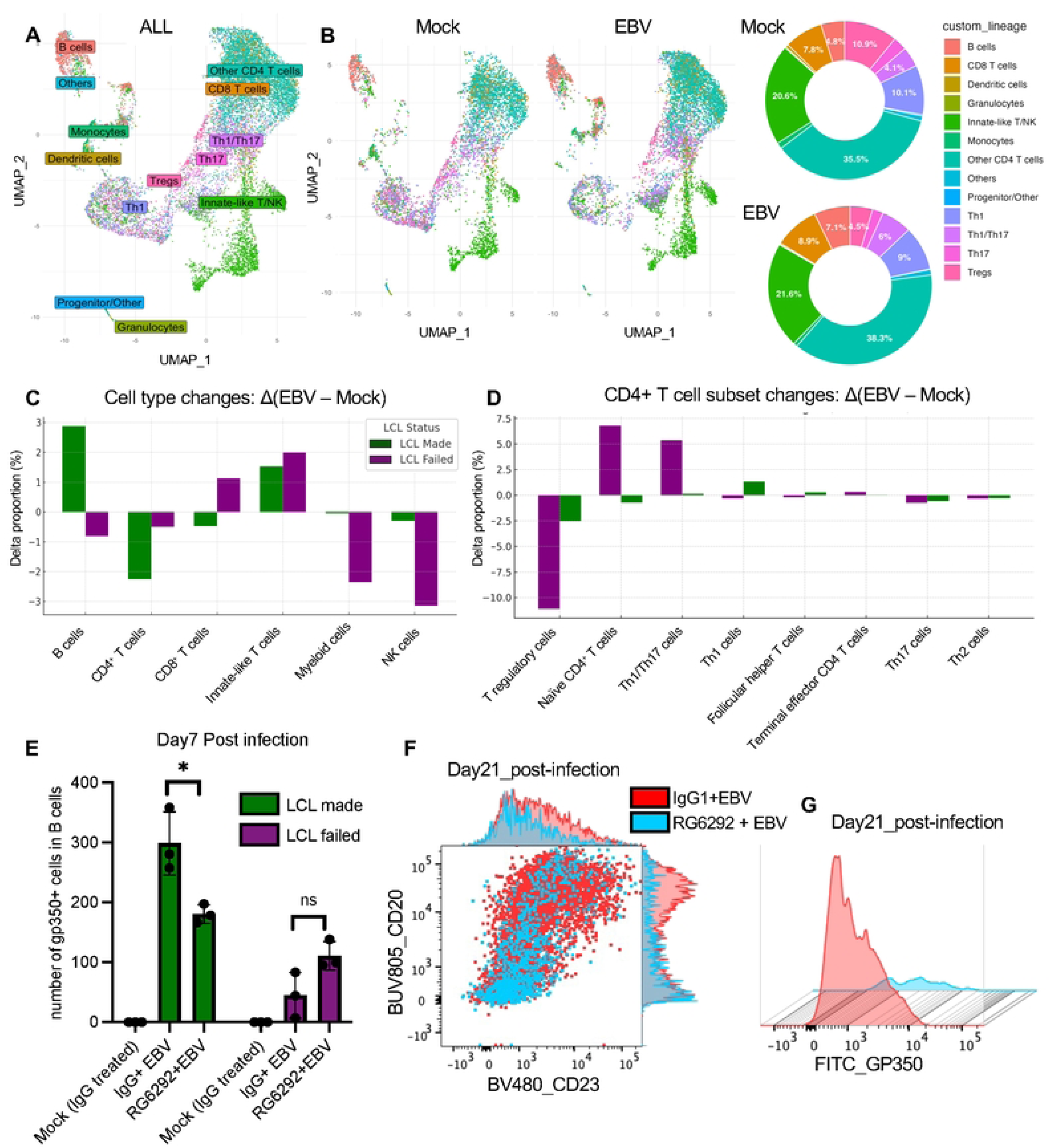
Single-cell RNA-seq and Treg depletion reveal regulatory control of EBV infection outcome. (A) UMAP of single-cell transcriptomes from all donors and conditions, annotated by major immune cell types. (B) UMAP plots and pie charts showing relative frequencies of immune lineages across mock and EBV-infected conditions. (C) Bar plot showing the change in cell type proportions (Δ = EBV – Mock) for each donor, grouped by LCL outcome. (D) Subset analysis of CD4⁺ T cells showing relative frequencies of Treg and naive CD4⁺ cells per donor and condition. (E) Quantification of gp350⁺ B cells at Day 7 post-infection in LCL-made and LCL-failed donors PBMCs treated with IgG1 isotype control (Mock), EBV + IgG1 isotype control or EBV + RG6292. Bars represent the number of gp350⁺ cells within the CD19⁺ B cell compartment (mean ± SEM; p < 0.05, ns = not significant; a paired two-sided Student’s t-test (n=3)). (F) Representative flow cytometry plot of CD23 and CD20 expression on CD19⁺ B cells at Day 21 post-infection. Red and light blue represent EBV-infected cultures treated with isotype control or RG6292, respectively. Marginal histograms show marker intensity distribution. (G) gp350 surface expression on CD19⁺ B cells at Day 21 post-infection. Histogram overlay compares isotype-(red) and RG6292-treated (light blue) conditions.

Within CD4⁺ T cells, Treg frequency decreased markedly in LCL-failed donors but was relatively stable in LCL-made donors (**Figure 6D**). IL2RA expression within Treg clusters was also reduced in LCL-failed but not in LCL-made donors (**Supplementary Figure S9 I and J**), consistent with bulk RNA-seq data (**Figure 5G**).

To test the functional role of Tregs in EBV infection control, we treated PBMCs with RG6292, a Fc-engineered anti-CD25 antibody that depletes Tregs via ADCC while preserving IL-2 signaling to effector cells [45, 46]. gp350 surface expression was used as a proxy for EBV-infected B cells. RG6292 treatment selectively reduced Tregs (CD4⁺ CD25⁺ FOXP3⁺) (**Supplementary Figure S9 A-H**). In LCL-made donors, RG6292 treatment reduced gp350⁺ B cells at Day7 post-infection, indicating that Treg depletion impairs EBV-driven B cell transformation (**Figure 6E**). In LCL-failed donors, where Treg-mediated regulation was already compromised, RG6292 had minimal additional effect. By Day 21, isotype control–treated, EBV-infected cultures showed B cell expansion and increased CD23 and gp350 expression, whereas RG6292-treated, EBV-infected cultures did not show B cell expansion or EBV infection marker expression, instead exhibiting extensive cell death, similar to that seen in LCL-failed donors (**Figure 6F-G**). These results support a model in which the Treg population facilitates EBV-mediated transformation, while Treg depletion mimics effective antiviral immunity.

### EBV Gene Programs Correlating with Infection Outcomes

We next set out to determine whether EBV gene expression correlated with infection outcomes (**Figure 7**). Bulk RNA-seq of PBMC did not yield significant EBV reads at the earliest time points after infection. To improve EBV transcript detection in RNA-seq, we used custom viral enrichment probes. Probe capture increased the percentage of viral reads by approximately 9.9-fold (paired Student’s t-test, p = 4.3e-09), with consistent enrichment across donors for each timepoint (**Supplementary Figure S10**). This allowed detection of low-abundance viral transcripts that were not observed in bulk RNA-seq of primary EBV infection. We analyzed EBV-infected PBMCs at Day 0 (preinfected), Day 1 and Day 7 post-infection (**Figure 7A**). Based on temporal dynamics, EBV genes clustered into three major groups (**Figure 7B**). The first group showed low expression at Day 0 (possibly detecting pre-infected B-cells), sharp induction at Day 1, and sustained high levels through Day 7. This group included EBNA-LP, BMRF1, and BFRF3, consistent with abortive lytic activation. The second group showed transient expression, with peak expression at Day 1 followed by a significant decrease by Day 7. These genes, such as BZLF1, BLLF1, and BGLF2, align with early lytic and replication-associated programs. The third group showed delayed expression, with minimal activity at early time points and increased expression by Day 7. This cluster included LMP1, LMP2B, and BNLF2a, which function in latency establishment and immune modulation. We next analyzed EBV gene expression patterns based on LCL outcome using previously characterized donor samples (Figure 7C). At Day 1, lytic genes such as BZLF1, BALF5, and BRRF1 showed higher expression in LCL-failed donors. In contrast, several latent and immune modulatory genes, including LMP1 and BCRF1, were more highly expressed in LCL-made donors. At Day 7, some lytic genes such as LF3 remained elevated in LCL-failed donors, while expression of latency-associated genes such as BCRF1, LMP2B, and BILF1 remained higher in the LCL-made group. These patterns, supported by individual gene trajectories (**Figure 7D**), indicate that LCL-failed donors exhibit elevated early lytic activation without proper transition to latency, which may compromise B cell transformation and long-term EBV persistence.

**Figure 7.**
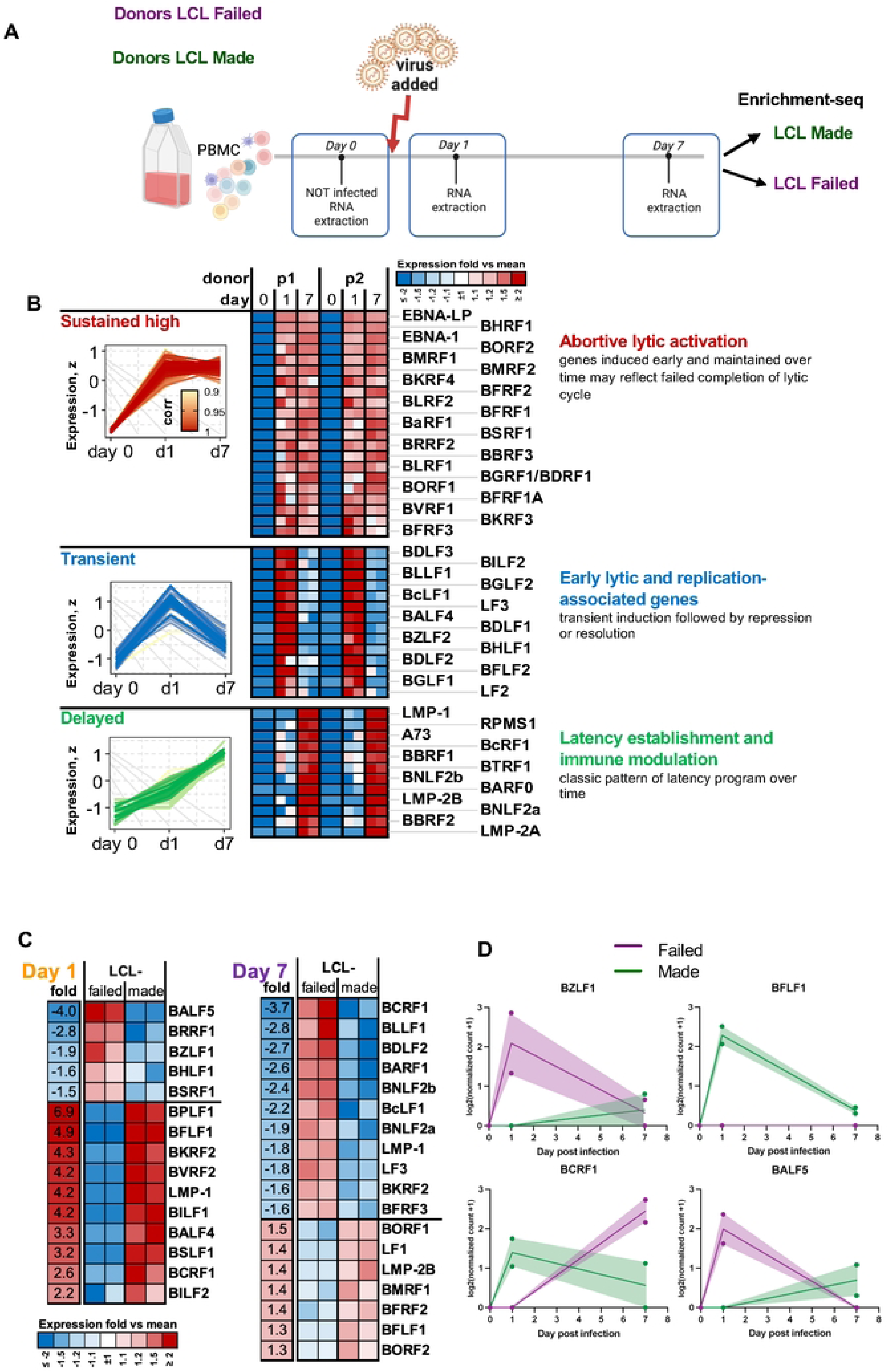
EBV Gene Expression Programs Diverge by Transformation Outcome in Infected PBMCs. (A)Schematic of *ex vivo* PBMC infection and EBV transcript enrichment-Seq workflow. PBMCs were infected with the MUTU1 strain of EBV and collected at Day 0, Day 1, and Day 7. Donors classified as LCL made or LCL failed were selected based on outcomes from a prior trial (see Supplementary Table 2); n=2. EBV-specific hybrid capture probes were used to increase viral transcript detection. (B) Temporal clustering of EBV genes based on expression patterns identified three major profiles: early and sustained (e.g., EBNA-LP, BMRF1), transient (e.g., LF3, BLLF1), and delayed (e.g., LMP1, BNLF2a). (C) Comparative EBV gene expression by LCL outcome across time points. LCL-failed donors showed higher early induction of selected viral genes, but lower expression of transcripts observed in LCL-made donors at Day 7. (D) Representative EBV genes demonstrate divergent expression trajectories between LCL-made (green) and LCL-failed (purple) donors. BZLF1 and BALF5 show elevated early expression in LCL-failed donors, while BFLF1 and BCRF1 are preferentially sustained or induced in LCL-made donors by Day 7.

## Discussion

In this study, we have used *ex vivo* EBV infection of PBMC to demonstrate a diversity of immune cellular and transcriptional responses among individuals that can lead to different outcomes with respect to EBV transformation of B-lymphocytes and LCL formation. We observed that LCL formation from PBMC is strictly donor dependent and highly reproducible within donor PBMCs. In contrast, purified B-cells from all donors formed LCLs with equally high success rates. Thus, donor variation in non-B cell components determine the outcomes of EBV infection and transformed B-cell outgrowths. Flow cytometry, bulk RNAseq and single cell RNA-seq revealed that at least two major phases of immune response could be easily characterized. An early response directed by monocytes, and a later response determined by a balance between Treg and CD8+ T cell activity. Overall, the *ex vivo* PBMC-based model provides a robust model to study these early cellular and transcriptional response to EBV infection.

The earliest transcriptional response to EBV was dominated by interferon signaling, oxidative stress responses, and lipid metabolic remodeling. Key regulators such as IRF1 and EIF2AK2 (PKR) were upregulated within 24 hours, suggesting a coordinated antiviral and stress response network. Lysosomal acid lipase (LIPA) emerged as a monocyte-specific antiviral factor, and its pharmacological inhibition increased viral load, which indicates a functional role in early EBV restriction. In parallel, chemokine receptor remodeling in monocytes, including reduced CCR1 and CCR2 expression, suggests that altered migratory or differentiation states may influence early immune control. These early changes define a rapid innate immune sensing phase that may shape downstream transformation potential and immune regulation.

The rapid early response to EBV infection involves a shift in monocyte lipid metabolism, marked by the upregulation of LIPA, which hydrolyzes cholesterol esters into free cholesterol and fatty acids. These lipids serve as substrates for oxysterol production via both enzymatic (e.g., CH25H) and non-enzymatic, ROS-mediated pathways [47, 48]. A recent study shows that LIPA is required for 25HC biosynthesis, linking lysosomal lipid hydrolysis to oxysterol production [49]. In addition, IRF1 activation at Day 1 can upregulate transcription of genes encoding NADPH oxidase components, which also elevate reactive oxygen species (ROS) in immune cells [50–52]. The resulting oxysterols, including 25-hydroxycholesterol (25HC) and 7α,25-dihydroxycholesterol, have been shown to bind to EBI2 (GPR183), a chemotactic GPCR that directs immune cell positioning during inflammation [53]. This oxysterol–EBI2 signaling axis has been found to produce 7α,25-OHC and induce macrophage migration in LPS-stimulated astrocytes [54–56]. 25HC has also been shown to directly inhibit both EBV and KSHV infection and gene expression [57], suggesting that oxysterols play dual roles in immune guidance and antiviral restriction. We propose that the LIPA–ROS–oxysterol–EBI2 axis is a key intersection between host lipid metabolism and EBV immune modulation (**Figure 8**).

**Figure 8.**
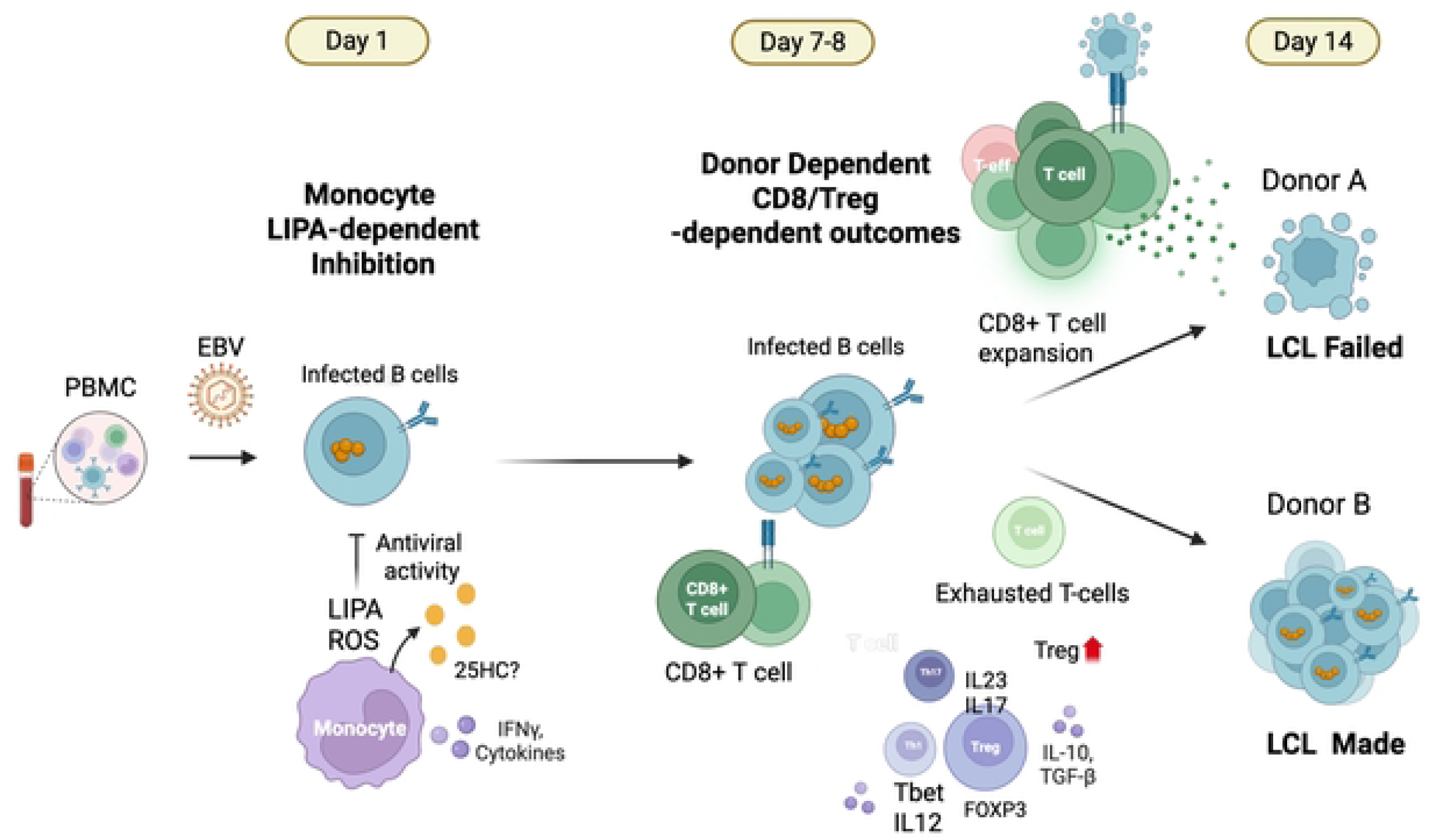
Summary model of the early and late phase immune responses to EBV infection of PBMC. Early response dominated by monocyte production of LIPA, with proposed pathways and cellular components related to LIPA expression, lipid metabolism, and EBV regulation. Later stage T-reg and CD8-T cell responses determine EBV infection outcome with respect to LCL-made or LCL-failed.

By Day 7, a secondary response to EBV infection of PBMCs was observed and correlated with donor-specific outcomes of LCL formation. Treg frequencies declined selectively in LCL-failed donors and remained stable in LCL-made donors, based on phenotypic analysis of CD4⁺CD25⁺FOXP3⁺ cells. These changes were mirrored by reduced IL2RA expression at the transcriptomic level, suggesting diminished Treg activity as a potential determinant of failed transformation. In contrast, increased Treg activity may foster a more permissive environment for viral persistence. The corresponding CD8⁺ T cell profiles further support this dichotomy. LCL-made donors exhibited a reduction in CD8⁺ T cells with increased expression of exhaustion-associated genes such as BATF, TOX, HAVCR2 (TIM-3), and ENTPD1 (CD39), which is consistent with the development of an exhausted CD8⁺ T cell phenotype. BATF, a transcription factor essential for chromatin remodeling and early differentiation of CD8⁺ T cells, and TOX, a master regulator of exhaustion, suggest that these cells are transitioning into a dysfunctional but persistent state. In contrast, LCL-failed donors showed elevated CD244, a marker of early CD8⁺ activation and pre-exhaustion, along with reduced expression of TOX and SLAMF6, indicating the absence of both terminally exhausted and progenitor-like CD8⁺ T cell subsets. This functional divergence shows that, CD8⁺ T cells may mount a more robust cytotoxic response that interferes with B cell transformation in LCL-failed donors whereas, regulatory and exhausted immune profiles allow EBV to establish latency and drive LCL formation in LCL-made donors.

Our data strongly suggest that Tregs can shape the immune environment required for EBV-driven B cell transformation. Flow cytometry, bulk and scRNA-seq indicate that Treg contraction occurred specifically in LCL failed outcomes, highlighting that early Treg contraction may reflect effective antiviral activity that restricts EBV-driven B cell transformation. To functionally test the role of Tregs, we treated PBMCs with RG6292, a non–IL-2–blocking anti-CD25 antibody that selectively depletes Tregs [45, 46, 58]. RG6292 treatment phenocopied the LCL-failed donor profile, suppressing gp350⁺ B cell emergence and abrogating EBV-driven transformation. A reduction in circulating CD4⁺CD25^+^ Tregs was previously reported in infectious mononucleosis patients, indicating that low Tregs accentuates the acute effector T-cell response to EBV [59]. This aligns with our findings that contraction of functional Tregs may reflect effective antiviral immunity, while their persistence supports EBV outgrowth. In addition, in nasopharyngeal carcinoma (NPC), higher tumor-infiltrating Tregs correlate with EBV DNA load and disease progression [60], suggesting that Treg presence may support EBV persistence and pathogenesis. Related findings from aging models show that impaired Treg function disrupts tissue homeostasis and repair in the CNS [61].

Treg regulation in EBV infection may be particularly relevant to autoimmune diseases such as multiple sclerosis (MS) where both EBV seropositivity and Treg dysfunction are implicated in disease pathogenesis [62, 63]. MS patients frequently show dysfunctional Tregs with impaired suppressive activity, and recent studies in a humanized mouse model demonstrated that EBV infection enhances effector T cell expansion and disrupts Treg homeostasis, correlating with increased CNS autoimmunity [62]. Mechanistically, inflammatory cytokines such as IL-6 and IL-1β have been shown to destabilize FOXP3 expression and reprogram Treg cells into dysfunctional states, suggesting that EBV-induced cytokine shifts could impair regulatory function and promote transformation-favorable environments [64]. MS patients more readily generate spontaneous LCLs (SLCLs) from PBMCs, even in the absence of exogenous virus, which likely reflects baseline immune dysregulation, including potential deficits in Tregs [65]. Treg dysfunction is a well-established feature across autoimmune diseases and therapeutic strategies such as low-dose IL-2 have been shown promising in selectively expanding functional Tregs and restoring immune tolerance in early-phase clinical trials [66, 67]. Our PBMC-based model captures donor-specific Treg dynamics and provides a human-relevant system to study how early regulatory imbalances influence EBV infection outcomes. These findings underscore the translational potential of this platform to dissect immunoregulatory circuits linking viral persistence, immune escape, and autoimmune risk.

At the viral level, EBV transcript enrichment-Seq revealed distinct gene expression trajectories during early infection. Based on temporal patterns, viral genes segregated into three groups: sustained expression (EBNA-LP, BMRF1), transient activation (BZLF1, BLLF1), and delayed induction (LMP2B, BNLF2a). Stratification by LCL outcome showed that LCL-failed donors exhibited higher expression of early lytic genes such as BZLF1 and BRRF1, while latent and immune modulatory genes including BCRF1 and LMP2B were elevated in LCL-made donors. These data suggest that successful transformation requires a temporal transition from lytic (or pre-latent) to latent gene expression, and that failure to resolve early lytic programs may compromise viral persistence and B cell immortalization. Transition to type III latency with sustained expression of LMP1 and LMP2, and full activation of the NF-kB activation pathways, have also been shown to correlate with LCL success in scRNAseq studies with enriched B-cell populations [17].

There are several limitations to our study. While the ex vivo infection of PBMC enables controlled study of early EBV–host responses, it lacks the spatial and multicellular complexity of lymphoid tissue. Bulk transcriptomic and flow cytometry analyses capture population-level dynamics but do not resolve cell–cell interactions or tissue-specific signaling cues. Future studies using spatial or single-cell approaches may help define how EBV reprograms immune coordination in physiologic settings. Our study is also limited using 8 donors, while a more expansive cohort might provide greater statistical power. Nonetheless, this model provides a tractable system to dissect temporal transcriptional and metabolic changes in primary human immune cells during the earliest stages of viral sensing.

In conclusion, the PBMC infection model provides a robust and tractable system to study EBV infection and immune response across different population cohorts. While we have focused here on a small collection of healthy donors, we were able to identify variations that correlate with infection outcomes. Future studies will expand the cohort size and the examination of populations with known or expected vulnerabilities to EBV infection.

## Material and methods

### Isolation and Infection of PBMCs

Peripheral blood mononuclear cells (PBMCs) were isolated from donor whole blood using density gradient centrifugation with Lymphoprep™ and SepMate™-50 tubes (Cat# 85460, STEMCELL Technologies, Vancouver, BC, Canada), following a modified manufacturer protocol. Whole blood was mixed 1:1 with DPBS containing 2% FBS, layered over 15 mL of Lymphoprep in SepMate tubes, and centrifuged at 1200 × g for 10 minutes at room temperature with full brake. The mononuclear cell layer was collected, diluted with DPBS/2% FBS to 50 mL, and centrifuged at 300 × g for 8 minutes. This wash step was repeated three times. PBMCs were then counted, pelleted, and cryopreserved in CryoStor® CS10 (Cat# 210202, BioLife Solutions Inc.) at a concentration of 2 × 10⁷ cells per vial.

For viral infections, cryopreserved PBMCs were thawed into RPMI medium (Cat# 11875085, Gibco) supplemented with 10% FBS, 1% GlutaMAX(Cat# 35050061, Gibco), and 100 U/mL penicillin, 100 µg/mL streptomycin (Cat# 15140122, Gibco), rested overnight, and plated at 1 × 10⁶ cells/mL in 12-well plates. Infections was performed using the Mutu1 viral strain at a multiplicity of infection (MOI) of 1. LCL-made samples were defined as by having a sustained increase in gp350⁺CD23⁺ cells over days 7-21, while LCL-failed donors showed an initial rise by Day 7 followed by a sharp decline by day 21.

### Virus production

Primary EBV infection experiments used virus derived from the Mutu I cell line, a Burkitt’s lymphoma-derived, EBV-positive line. Lytic reactivation was induced using 2 μM sodium butyrate and 20 ng/mL TPA for 72 hours. Following induction, both cells and culture medium were collected and to centrifuged twice at 3000 rpm for 10 minutes to eliminate cellular debris. The cleared supernatant was passed through a 0.45 μm filter and virus was concentrated by ultracentrifugation through a 10% sucrose cushion at 27,000 × g for 90 minutes at 4 °C. Viral pellets were resuspended in RPMI medium and the viral load quantified using droplet digital PCR (ddPCR).

### ddPCR

Viral DNA quantification was performed using ddPCR. A total of 8 µL of resuspended virus was first treated with DNase I (Cat# EN0521, Thermo Scientific™) to eliminate extracellular host genomic DNA, followed by incubation with Proteinase K (final concentration 20 μg/mL) at 56 °C for 30 minutes to inactivate DNase and digest nucleic acid–protein complexes. After enzymatic treatment, the enzymatic reaction was inactivated by heating at 95 °C for 10 minutes, then cooled to room temperature. Viral DNA was then used directly or diluted 1:2.5 with nuclease-free water ddPCR analysis. Reactions were prepared following the manufacturer’s protocol and quantified on the QX200 Droplet Digital PCR system (Bio-Rad). For the duplex ddPCR, Ribonuclease P protein subunit 30 (RPP30) was used as the housekeeping gene with primers and a VIC/MGB probe to quantify host genomic input[68]. LMP1 and BamHI-W regions were targeted using primers and a FAM/MGB probe designed for ddPCR using NCBI Primer Blast and Primer3Plus (Forward: AAGGTCAAAGAACAAGGCCAG; Reverse: GCATCGGGAGTCGGTGG; Probe: 6FAM–AGCGTGTCCCGTGGAGG–MGBNFQ)[69].

### Extracellular staining for cell surface markers and intracellular staining

For flow cytometric analysis, PBMCs were stained using three separate antibody panels tailored for characterizing lymphocyte subsets, monocytes, Tregs and EBV-infected cells. Viability staining was first performed using either Zombie-NIR (Cat# 423105, BioLegend) or LIVE/DEAD™ Fixable Aqua (Cat # L34966, Thermo Fisher Scientific), with incubation for 20 minutes at room temperature. To reduce nonspecific antibody binding, cells were then treated with Human TruStain FcX™ Fc receptor blocking reagent (BioLegend) and incubated for 20 minutes on ice.

Surface staining was performed in FACS buffer (PBS containing 5% FBS and 0.1% sodium azide) for 30 minutes at 4 °C with a cocktail of fluorochrome-conjugated monoclonal antibodies. Antibodies used included CD3_BB700 (clone SK7, BD Biosciences), CD4_ PE-Cy7 (clone SK3, BioLegend), CD4_Alexa Fluor 700 (clone OKT4, BioLegend), CD8_PE-Cy5 (clone SK1, BioLegend), CD25_BB700 (clone BC96, BD Biosciences), FOXP3_Alexa Fluor 647 (clone 259D, BioLegend), Live/Dead Fixable Aqua (Thermo Fisher Scientific), CD19_APC-H7 (clone SJ25C1, BD Biosciences), CD56_BV605 (clone NCAM16.2, BD Biosciences), CD16_BV711 (clone 3G8, BD Biosciences), CD20_BUV805 (clone 2H7, BD Biosciences), CD23_BV480 (clone M-L233, BD Biosciences), GP350_FITC (clone C61H, Invitrogen), CD14_BUV395 (clone MφP9, BD Biosciences), CD11c_BUV661 (clone B-ly6, BD Biosciences), CD64_APC (clone S18012C, BioLegend), HLA-DR_PE-Cy7 (clone L243, BioLegend), CCR1_PE (clone 5F10B29, BioLegend), CCR2_APC-Fire (CD192, clone K036C2, BioLegend), LIPA_DyLight488 (custom-conjugated), GP350_DyLight350 (custom-conjugated). Panel-specific combinations were optimized based on fluorochrome compatibility and marker co-expression patterns.

After surface staining, cells were washed twice with FACS buffer and resuspended in 400 μL of FACS buffer for acquisition. For FOXP3 staining, intracellular fixation and permeabilization were performed using the Intracellular Fixation & Permeabilization Buffer Set (eBioscience, Thermo Fisher Scientific), following the manufacturer’s protocol.

Flow cytometry was performed on a BD FACSymphony A3 (BD Biosciences). Compensation was calculated using UltraComp Beads (BD Biosciences) and single-stained controls. Gating strategies included doublet exclusion (FSC-A vs. FSC-H), viability gating, and lineage identification based on the antibody panels described above. Tregs were identified as CD3⁺CD4⁺CD25⁺FOXP3⁺ cells. NK cells were classified based on CD56 and CD16 expression. Monocytes were identified as CD14⁺CD64⁺HLA-DR⁺, and EBV-infected populations were assessed by GP350 and CD23 staining. Data were analyzed using FlowJo software (v10, BD Biosciences). Flow cytometry gating strategies are shown in Supplemental Figures S2, S2, S9).

### Lalistat2 treatment

PBMCs were pre-treated with 30 μM Lalistat2 (Cat# 1234569-09-5, Sigma-Aldrich) or an equivalent volume of DMSO for 30 minutes at 37°C prior to EBV infection [70]. Cells were then infected with the MUTU1 strain of EBV under standard culture conditions and maintained in the presence of Lalistat2 for the duration of the assay.

### RG6292 Treatment

Vopikitug (RG6292; MedChemExpress, Cat# HY-P990086) was used at 800 ng/mL, diluted in RPMI cell culture medium [58]. An equivalent concentration of human IgG1 isotype control (MedChemExpress, Cat# HY-P99001) was used in control conditions (Mock and EBV). Cells were then infected with the MUTU1 strain of EBV under standard culture conditions.

### Bulk RNA-seq and Analysis

Mock and EBV infected samples were collected at different time-points. Total RNA was isolated from 1 × 10^6^ cells using the RNeasy Mini Kit (Cat# 74104, Qiagen) followed by on-column DNase treatment per the manufacturer’s protocol. RNA quality was assessed at the Wistar Institute Genomics Core, with all samples showing RIN values >7 (Agilent TapeStation). Libraries were prepared using the QuantSeq 3’ mRNA-Seq kit (Lexogen), following standard protocols for generation of Illumina-compatible libraries. Sequencing was performed on an Illumina NextSeq 2000 using high-output mode, generating approximately 5 million single-end 75 bp reads per sample.

RNA-seq data were aligned using the STAR [71] algorithm to a combined reference of the human genome (hg19) and the EBV genome (NC_007605.1). RSEM v.1.2.12[72] software was used to estimate read counts and reads per kilobase per million mapped reads (RPKM) values using gene information from Ensemble transcriptome (v.GRCh37.p13) [72]. DESeq2[73] was used on raw counts to estimate significance of expression differences between any two experimental groups and generate normalized counts. Heat maps were generated using the pheatmap R package, and additional data visualizations were created with ggplot2. Gene set enrichment analysis was performed using QIAGEN’s Ingenuity Pathway Analysis software (IPA, QIAGEN; www.qiagen.com/ingenuity) using the ‘Canonical pathways’ and ‘Upstream regulators’ option. Additional pathway analysis was performed using Enrich R [74].

### Single-cell RNA seq and analysis

Single-cell transcriptomic analysis was performed using the Parse Biosciences Evercode Whole Transcriptome mini kit (Parse Biosciences, Seattle, WA), following the manufacturer’s protocol. Briefly, cells were first fixed and barcoded using split-pool combinatorial indexing, enabling non-droplet-based single-cell RNA capture. Following barcoding, cDNA was amplified and sequencing libraries were prepared using the Parse library construction workflow. Libraries were sequenced on an Illumina NovaSeq 6000 platform to achieve a target depth of approximately 20,000 reads per cell. Processed FASTQ files were aligned to a custom GRCh38 human – EBV(NC_007605.1) reference and quantified using the split-pipe pipeline. The resulting data was loaded into R, and the Seurat package [75] was used to create a single-cell object (CreateSeuratObject()). The data was then demultiplexed by condition and patient sample ID. A standard processing workflow was applied (NormalizeData(), FindVariableFeatures(), and ScaleData()). Seurat was also used to identify differentially expressed genes (FindVariableFeatures()) as well as post-processing dimensional analysis (RunPCA(), FindNeighbors(), FindClusters(), and RunUMAP()). The Azimuth [76] package was used to identify cell types.

### EBV Transcript Enrichment-Seq by Hybrid-Capture

To enhance detection of EBV transcripts in RNA-seq libraries, hybrid-capture enrichment was performed using a custom panel of EBV-specific probes. Custom hybrid capture probes were designed and synthesized to enrich for EBV-specific sequences. Briefly, the unique sequences of the entire EBV genome according to both major strains (Type I NC_007605 and Type II NC_009334) were segmented into 120-bases and designated as non-repeat probe sequences with back-to-back tiling across the viral genome. The repetitive elements were identified and similarly segmented into 120-bases for the first ‘unit’ of each repeat element. Single-stranded DNA probes were synthesized using all the 120-base sequences by IDT into a non-repeat pool and a repeat pool (Integrated DNA Technologies, Coralville, IA). Prior to hybrid capture, these pools were combined into single hybrid capture probe pool with approximately 7.5 parts of the non-repeat probe pool and 2.5 parts of the repeat probe pool. Library preparation followed the standard Illumina TruSeq stranded Total RNA protocol, after which enriched libraries were generated using the IDT xGen Hybridization and Wash Kit (Integrated DNA Technologies) according to the manufacturer’s protocols. Enriched libraries were amplified and purified before sequencing on an Illumina platform.

## Data Availability

All RNA-seq data is deposited in NCBI GEO datasets under the project XXXX.

## Acknowledgements

We thank the Wistar Institute Shared Resources Facilities in Genomics, Bioinformatics, Flow Cytometry and Phlebotomy. This work is supported by grants from the DOD (MS220073) and NIH NCI P01 CA281867 and R01 AI153508 to PML, R50 CA211199 to AK, and U01 CA284811 to RFA and RRX.

## Supplementary Data

**Supplementary Table S1. Donor demographic and experimental metadata.**

Summary of donor characteristics used in EBV infection studies. The table includes donor ID, age, sex, race, and health status, along with experimental annotations indicating whether lymphoblastoid cell lines (LCLs) were successfully generated (Made, Failed, or M/F for mixed outcomes). Columns also denote availability of RNA-seq and flow cytometry data, as well as EBV serostatus.

**Supplementary Table S2. Summary of LCL outgrowth efficiency across donors and infection conditions.**

Donor PBMCs were infected with EBV at varying multiplicities of infection (MOIs) ranging from 0.1 to 10, and LCL outgrowth was assessed across 8–11 independent trials. Each cell represents the outcome of a single trial under the indicated condition. Donors highlighted in green successfully formed LCLs (LCL Made), while those in purple failed to establish LCLs (LCL Failed) under cyclosporin A–untreated conditions. “o” indicates successful LCL outgrowth; “x” indicates failure. Gray-shaded columns represent MOI conditions not tested for the corresponding donor.

**Supplementary Figure S1. Success rate of LCL outgrowth per donor.**

Bar plot shows the percentage of successful LCL outgrowths per donor under untreated (green) and Cyclosporin A–treated (gray) conditions. Each bar represents the proportion of successful EBV infections out of total attempts across multiple MOI conditions (see Supplementary Table 2).

**Supplementary Figure S2. Flow cytometry gating strategy for major immune cell subsets**

Cells were gated sequentially for lymphocyte population (FSC-A vs. SSC-A), single cells (FSC-A vs. FSC-H), and live cells (Aqua green Live/Dead dye). Lineage-specific markers were then applied within the live, singlet gate to define major immune subsets, including monocytes, T cells, NK cells, and B cells.

**Supplementary Figure S3. Gating strategy for identification of regulatory T cells (Tregs).**

Flow cytometry plots showing sequential gating to identify CD4⁺FOXP3⁺CD25⁺ regulatory T cells.

**Supplementary Figure S4. Consistent upregulation of HIF1A and KLRF1 following EBV infection.**

(A) Expression of HIF1A and (B) KLRF1 in PBMCs following *ex vivo* EBV infection, compared to mock-treated controls, across all timepoints (Days 1, 7, and 14). Boxplots show log2-transformed normalized gene counts. p-values were calculated using a paired Student’s t-test.

**Supplementary Figure S5. LIPA expression and potential mechanism of LIPA in EBV-infected cells.**

(A) Flow cytometry plots showing LIPA expression in CD19⁺ B cells (left), CD3⁺ T cells (middle), and CD14⁺ monocytes (right). (B) Relative LIPA mRNA expression measured by qPCR in PBMC infected with EBV compared to mock at Day1. Expression normalized to GUSB. ***p < 0.001 by paired t-test.

**Supplementary Figure S6. Unsupervised clustering of differentially expressed genes at Day 14.**

Heatmap showing hierarchical clustering of over 600 differentially expressed genes (rows) at Day 14, comparing donors in which LCLs were successfully established (green) or failed (purple). Scaled expression values are shown (Z-scores per gene). Clustering was performed using Euclidean distance and complete linkage.

**Supplementary Figure S7. Key immune surveillance and ROS-related genes show distinct expression in LCL made versus failed donors.**

(A) Heatmap of selected differentially expressed genes (rows) related to immune activation and redox signaling, comparing LCL made (green) and failed (purple) donor groups at Day 14. Expression values are scaled by gene (Z-score). (B-K) Boxplots showing log-transformed normalized expression values of representative genes in EBV-infected and mock conditions for each group.

**Supplementary Figure S8. Differential expression of T cell exhaustion-related genes in EBV-infected PBMCs from LCL-failed and LCL-made donors.**

Boxplots display log₂ normalized expression values (counts +1) for each gene in peripheral blood mononuclear cells (PBMCs), stratified by donor LCL outcome (Failed vs Made) and infection condition (Mock [blue] vs EBV [red]). (A) ENTPD1, (B) HAVCR2, (C) TOX, (D) SLAMF6.p-values indicate results of unpaired two-tailed Student’s t-tests between EBV and mock conditions within each donor group.

**Supplementary Figure S9. Treg Gating Strategy and Depletion by RG6292 Treatment**

(A–C) Flow cytometry gating strategy: lymphocytes were first identified by forward and side scatter (A), followed by singlet discrimination (B), and viability gating using Aqua green live/dead stain (C). (D) CD4⁺ and CD8⁺ T cells were gated from live lymphocytes. (E–F) Representative plots showing frequency of CD25⁺FOXP3⁺ Tregs within the CD4⁺ compartment in mock-IgG1 treated (E) and RG6292-treated (F) PBMCs. RG6292 selectively depleted CD4⁺CD25⁺FOXP3⁺ Tregs. (G–H) gp350 expression on CD19⁺ B cells at Day 7 post-infection in IgG + EBV-infected (G) or RG6292-treated EBV infected (H) cultures. (I–J) Single-cell RNA-seq violin plots showing IL2RA expression in Treg clusters from LCL-made (I) and LCL-failed (J) donors under mock and EBV conditions.

**Supplementary Figure S10. Enhanced EBV transcript detection with probe enrichment-Seq.**

(A) Heatmap of normalized EBV counts (log₂ scale) across Day 0, Day 1, and Day 7 post-infection samples. (B) Line plot comparing log₂(% EBV reads) in samples with and without EBV probe enrichment across multiple donors and time points. (C) Boxplot of log₂(% EBV reads) showing a statistically significant increase in EBV reads in the with probe condition compared to without probe (paired Student’s t-test, p = 4.3e-09).

